# Dynamic switching between intrinsic and extrinsic mode networks as demands change from passive to active processing

**DOI:** 10.1101/2020.03.18.997031

**Authors:** Frank Riemer, Renate Grüner, Justyna Beresniewicz, Katarzyna Kazimierczak, Lars Ersland, Kenneth Hugdahl

**Affiliations:** Mohn Medical Imaging and Visualization Center, University of Bergen and Haukeland University Hospital, Bergen, Norway; Department of Radiology, Haukeland University Hospital, Bergen, Norway; Department of Physics and Technology, University of Bergen, Norway; Department of Biological and Medical Psychology, University of Bergen, Norway; Department of Clinical Engineering, Haukeland University Hospital, Bergen, Norway; Division of Psychiatry, Haukeland University Hospital, Bergen, Norway

**Keywords:** fMRI, default mode network, extrinsic mode network, working memory, mental arithmetic, mental rotation

## Abstract

We here report on the relationship between default and extrinsic mode networks across alternating brief periods of rest and active task processing. We used three different visual tasks: mental rotation, working memory and mental arithmetic in a classic fMRI ON-OFF block design where task (ON) blocks alternated with equal periods of rest (OFF) blocks. By analysing data in two ways, using an ON-OFF contrast, we showed the existence of a generalized task-positive network, labelled the extrinsic mode network (EMN) which was anti-correlated with the default mode network (DMN) as processing demands shifted from rest to active processing. We then identified two key regions of interest (ROIs) in the SMA and Precuneus/PCC regions as hubs for the extrinsic and intrinsic networks, and extracted the time-course from these ROIs. The results showed a close to perfect correlations for the SMA and Precuneus/PCC time-courses for ON-respective OFF-blocks. We suggest the existence of two large-scale networks, an extrinsic mode network and an intrinsic mode network, respectively, which are up- and down-regulated as environmental demands change from active to passive processing.

## Introduction

Ever since the discovery and identification of the default mode network (DMN) by Marcus Raichle and colleagues^1–3^ the study of network interactions has been a major issue in imaging neuroscience^4^. This has in particular concerned intrinsic interactions and anti-correlations between the DMN, as a resting-state network, and suggested task-positive networks, which are activated during active task-processing^5–8^. Other studies have focused on the decomposition of the DMN into sub-networks typically activated in task-processing situations, such as the salience and executive networks^9–13^. At around the same time as the discovery of the DMN, Duncan and Owen^14^ suggested that regions in the dorsolateral and anterior cingulate frontal cortex were activated to multiple processing demands. Duncan and Owen^14^ described their discovery as a *“regional specialization of function within prefrontal cortex*” (p. 475), which generalized across a variety of cognitive tasks and situations. The initial discovery by Duncan and Owen^14^ were later replicated and extended by Fedorenko et al.^15^, who found a common network structure across seven memory, executive, and attention tasks, and by Duncan^16^ who now labelled these activations a multiple demand (MD) system. Adding to this, Hugdahl et al.^17^ analysed data from nine separate fMRI experiments and found a generalized task non-specific network activated across all nine tasks which involved the SMA/anterior cingulate, lateral prefrontal cortex, and inferior parietal lobule. Hugdahl et al.^17^ labelled this network, the extrinsic mode network (EMN) to distinguish it from the intrinsic DMN. It is therefore clear that not only is there a set of network activations during resting periods, which collectively could be called non-specific task-negative networks, but also a set of similar non-specific task-positive networks, that has been variously labelled multiple demand system^16^, frontal-lobe network^14^, or extrinsic mode network^17^. We will use the term “extrinsic mode network (EMN)” in the following to describe this generalized task-positive network. An important question is how the DMN and EMN networks interact with regard to dominant up- and down-regulations across time when environmental demands repeatedly change from active to passive task-processing, i,e, change from engagement to rest and vice versa, on a short-term basis. Most studies of the relationship between task-negative and task-positive networks are either conducted during prolonged resting-periods, or having subjects solve a single task, addressing a single cognitive domain^1,10,18^, like working memory or attention as examples. This would however not capture the question of the dynamics of resting-state and non-specific task-positive network interactions, where tasks change from one processing period to another across the experimental session. Hugdahl et al.^19^ suggested an experimental proxy to the everyday switching between periods of task engagement alternated with periods of rest, where tasks moreover will differ in terms of cognitive domain and processing load from one processing period to another. In a fMRI situation, this can be obtained by alternating task-presence and task-absence periods, using a traditional fMRI block-design^20^, with ON- and OFF-blocks representing, active versus passive processing periods, respectively. Such an approach was taken by Hugdahl et al.^19^ who used an auditory dichotic listening task^21–23^ with ON-blocks with pseudo-random presentations of three different cognitive tasks involving perception, attention, and executive function that were interspersed in between OFF-blocks with no tasks present. The results showed statistically significant anti-correlations between the DMN and EMN, particularly in the inferior frontal and posterior cingulate cortex regions. Moreover, EMN up-regulations at the transition from an OFF- to an ON-block were steeper and more prolonged, than the corresponding up- regulation of the DMN in the transition from an ON- to an OFF-block. The study by Hugdahl et al.^19^ was however confined to the auditory modality, and had the different cognitive domains embedded within a single task, the so called “forced-attention dichotic listening task”^22^. Thus, it is not known if a similar pattern of interactions between the DMN and EMN would hold for; a) the visual modality, b) when splitting the cognitive domains across tasks, and c) expanding the tasks to more complex cognition, like number arithmetic, working memory, and mental rotation. The aim of the study was therefore to provide an extended replication of the Hugdahl et al.^19^ study by including different tasks, and processing strategies, while staying within the same basic experimental approach as the one used by Hugdahl et al.^19^. The three tasks were chosen because they represent three cognitive domains typically encountered during an ordinary day, visuo-spatial processing, mental arithmetic, and demands for handling information in short-term memory for brief periods of time. A mental rotation task^24^ was chosen because this task is shown to provide significant fronto-parietal activations^25–27^. A working memory task was chosen because working memory is a central cognitive concept, which includes attention and executive control in addition to short-term memory ^28,29^, and would thus be a valid proxy for the varying processing demands during an ordinary work-day in life. Arithmetics with adding numbers was chosen because it draws on a common cognitive ability encountered every day, the ability for mental arithmetic and to manipulate numbers. Previous research has shown that this task in isolation produces reliable activation in inferior frontal cortex and anterior cingulate^30,31^. In addition to whole brain analysis, we chose two region-of-interests (ROIs) for comparisons and correlations between the DMN and EMN networks, respectively. For the DMN we chose the precuneus as ROI, since this region is consistently activated during DMN up-regulations^5,32^. For the EMN we chose the intersection of anterior cingulate (ACC) and supplementary motor area (SMA) since this region is consistently activated during periods of EMN up-regulation^15,17^. The aim of the present study was therefore to replicate and extend our previous study^19^ on the dynamic interaction and correlation between the DMN and EMN cortical networks. Considering the current replication crisis in neuroscience and psychology^33^, an extended replication is a necessary first step for establishing a solid factual basis for new findings.

## Results

### Behavioural data

Mean response accuracy was calculated as the ratio of correct responses to overall number of responses and expressed as a hits ratio. For the mental rotation task, the hits ratio was 0.560, for the working memory task it was, and 0.769, and for the mental arithmetic task it was 0.843. The corresponding Cohen’s d values were; 0.55, 0.91, and 1.84, for the mental rotation, working memory, and mental arithmetic tasks, respectively. Following a standard interpretation of corresponding effect sizes, all three tasks showed medium to strong effect size. Mean response latency, or reaction-time (RT), for the three tasks were 404.2 ms (SD 143.0) for the mental rotation task, 168.4 ms (SD 56.5) for the working memory task, and 218.3 ms (SD 38.0) for the mental arithmetic task. These results confirm that the subjects had understood and performed the tasks as expected.

#### fMRI data

Figure 1 shows the results from the inclusive conjunction analysis of mean joint activations across the three tasks, *p* < 0.05, FWE-corrected, with significant activations for the ON-OFF contrast seen in the left-hand panel, and corresponding activations for the OFF-ON contrast seen in the right-hand panel.

**Figure 1.**
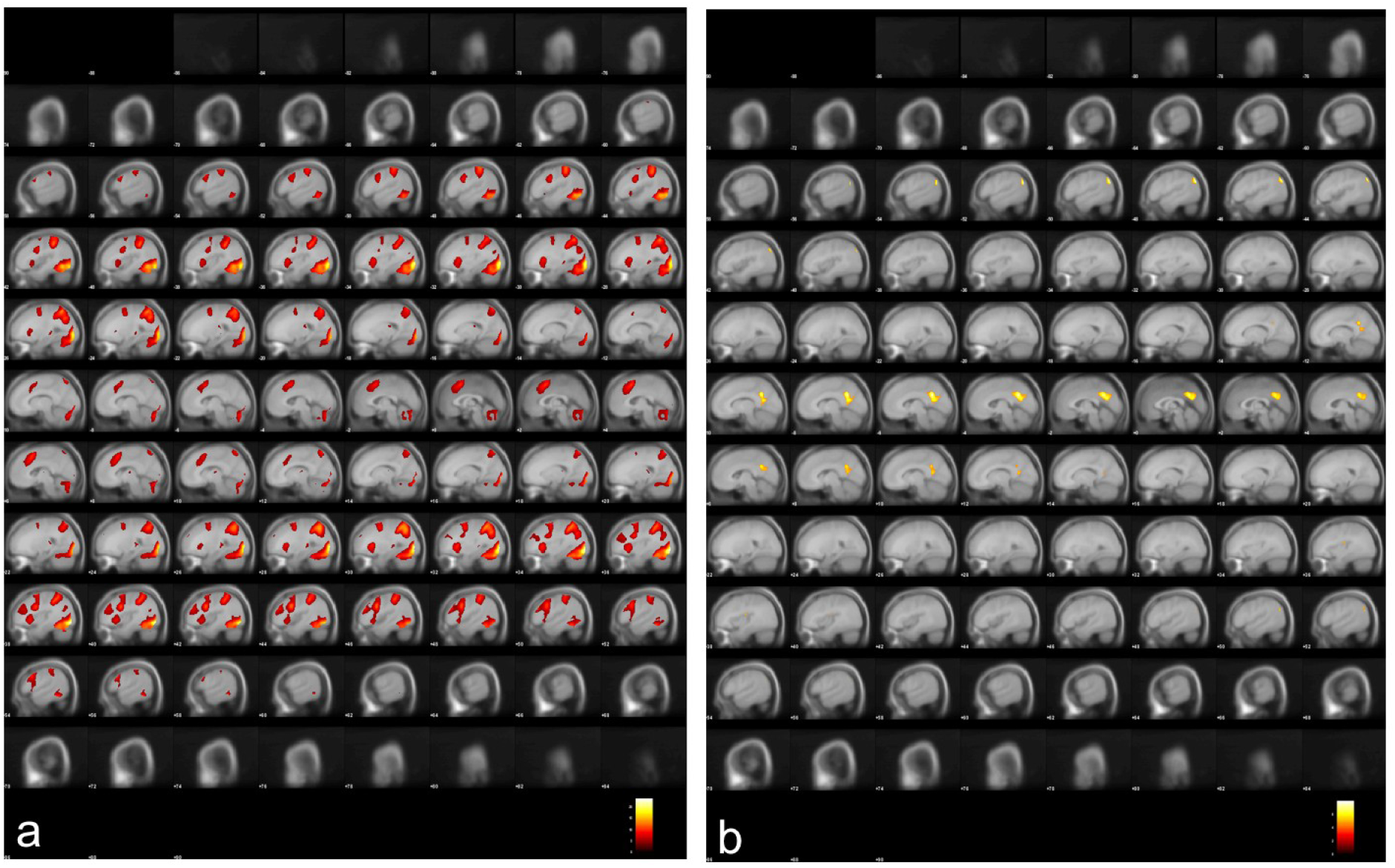
Activations that passed the .05 FWE-corrected significant threshold, activation ROI min. 20 voxels, shown in 2 mm sagittal slices through the whole brain volume. The panel (**a**) to the left shows activations obtained during ON-blocks contrasted with activations obtained during OFF-blocks (ON – OFF). The panel (**b**) to the right shows activations obtained with the contrast flipped, i.e. obtained during OFF-blocks contrasted with activations obtained during ON-blocks (OFF – ON). See results for further details.

The ON-OFF contrast produced significant, *p* = 0.05 FWE-corrected, activations in the right SMA (MNI coordinates: x 4, y 14, z 50), right inferior occipital gyrus (x 32 y −85 z −32) right angular gyrus (x 30, y −64 z44); left superior parietal lobule (x −26, y −58, z 54); right and left precentral gyrus (x 46, y 10, z 32, and x −46, y 4 z 30, respectively), left anterior insula (x −30, y 18, z 6), left middle frontal gyrus (x −26, y −4, z 50). All t-values > 7.46, critical .05 t-value = 4.40 with 447 df. The OFF-ON contrast produced corresponding activations in the left precuneus/posterior cingulate gyrus (x −6, y-54, z 32/ x 0, y 52 z 31), left and right angular gyrus (x −48, y −70, z 36 and x 50 y −68, z 30), and in the right central operculum (x 38, y −14, z 18). All t-values > 4.97, critical .06 t-value = 4.40. Figure 2 shows the results split for the three tasks, with activations overlaid on the MRICron anatomical template (ch2.better.niftii) as different layers and colours, see text in Figure legend for explanation of colour codings. As seen in Figure 2, the main findings from the overall conjunction analysis were confirmed in the separate analyses, with similar patterns of activations for all three tasks.

**Figure 2.**
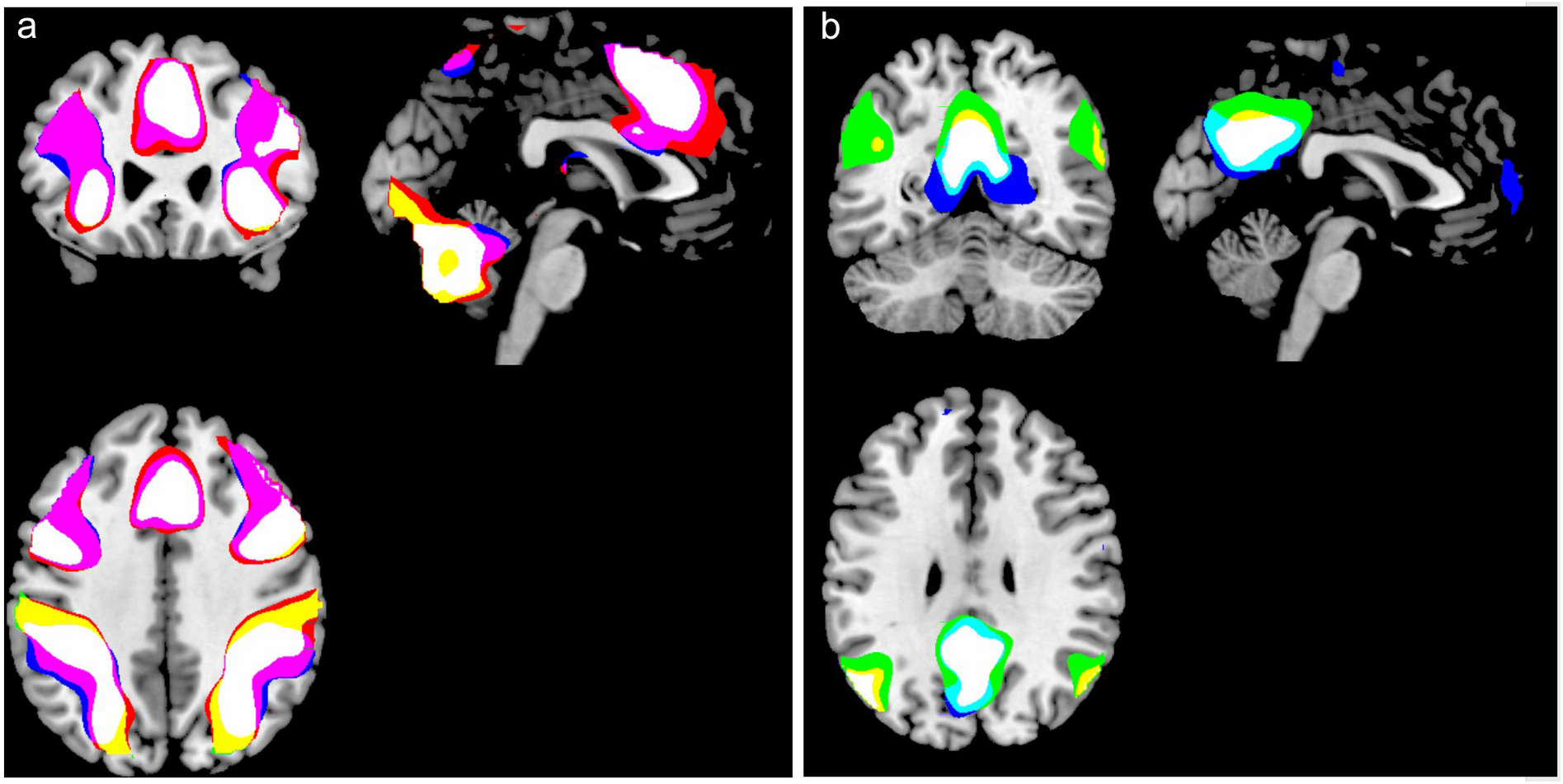
Activations for the three separate tasks overlaid on top of each other, with activation caused by the working memory task in red, by the mental rotation task in blue and by the mental arithmetic task in green. The panel (**a**) to the left shows activations obtained during ON-blocks contrasted with activations obtained during OFF-blocks (ON – OFF). The panel (**b**) to the right shows activations obtained with the contrast flipped, i.e. obtained during OFF-blocks contrasted with activations obtained during ON-blocks (OFF – ON). Common activations across all three tasks are correspondingly indicated in white colour. Common activations for the mental rotation and working memory tasks are indicated in violet colour. Common activations for the mental rotation and mental arithmetic tasks are indicated in cyan colour, and common activations for the working memory and mental arithmetic tasks are indicated in yellow colour. There is substantial overlap in the activation patterns across the three task. By visual inspection, mental rotation and working memory overlap the most (ON-OFF), while the mental arithmetic task provides the largest extent of activation for the contrast (OFF-ON).

Analysing the contrasts separately for each task, also yielded activations beyond what was seen in the conjunction analysis across tasks. For the ON-OFF contrast, the Mental rotation task yielded additional activation in the right thalamus (x 24, y −30, z 4) and the Mental arithmetic yielded additional activations in right and left thalamus (x 26, y −30, z 2, and x −24, y −30, z 0, respectively). For the OFF-ON contrast, the corresponding additional activations for the Working memory task were seen in the left and right lingual gyrus (x −24 y −44, z −8 and x 30, y −38, z −12, respectively), in the left medial and lateral superior frontal gyrus (x −2, y 58, z 8 and x −16, y 38, z 52, respectively), and in the right posterior insula (x 38, y −12, z 16). For the Mental arithmetic task, the corresponding additional activations were seen in the right cerebellum (x 30, y −74, z −40), left middle temporal gyrus (x −62, y 34, z 48), and in the left superior frontal gyrus (x −22, y 34, z 48).

We next identified overlapping activations for all three tasks in the SMA and precuneus/PCC, as regions-of-interest (ROIs), as these regions are key nodes in the EMN and DMN networks, respectively^2,17^. These regions were thereafter overlaid on an anatomy template, which is shown in the upper panel of Figure 3a-b. We then extracted the mean time-course from these regions, separately for ON- and OFF-blocks across the scanning session, which is seen in the lower panel of Figure 3c. Windowed correlation coefficients were calculated with a time-window of 34 sec (the length of one ON- or OFF-block. Development of correlation coefficients across time is shown as an intermittent black line in Figure 3c. As seen in the Figure, there was a strong negative correlation at the beginning and end of each task-related ON-block, mean Pearson r-coefficients at points of inflection was –0.56 ± 0.21. A corresponding strong positive correlation was seen at approximately the middle of each task ON- and rest OFF-periods, mean peak Pearson R's was +0.68 ± 0.19. See intermittent line in Figure 3c.

**Figure 3.**
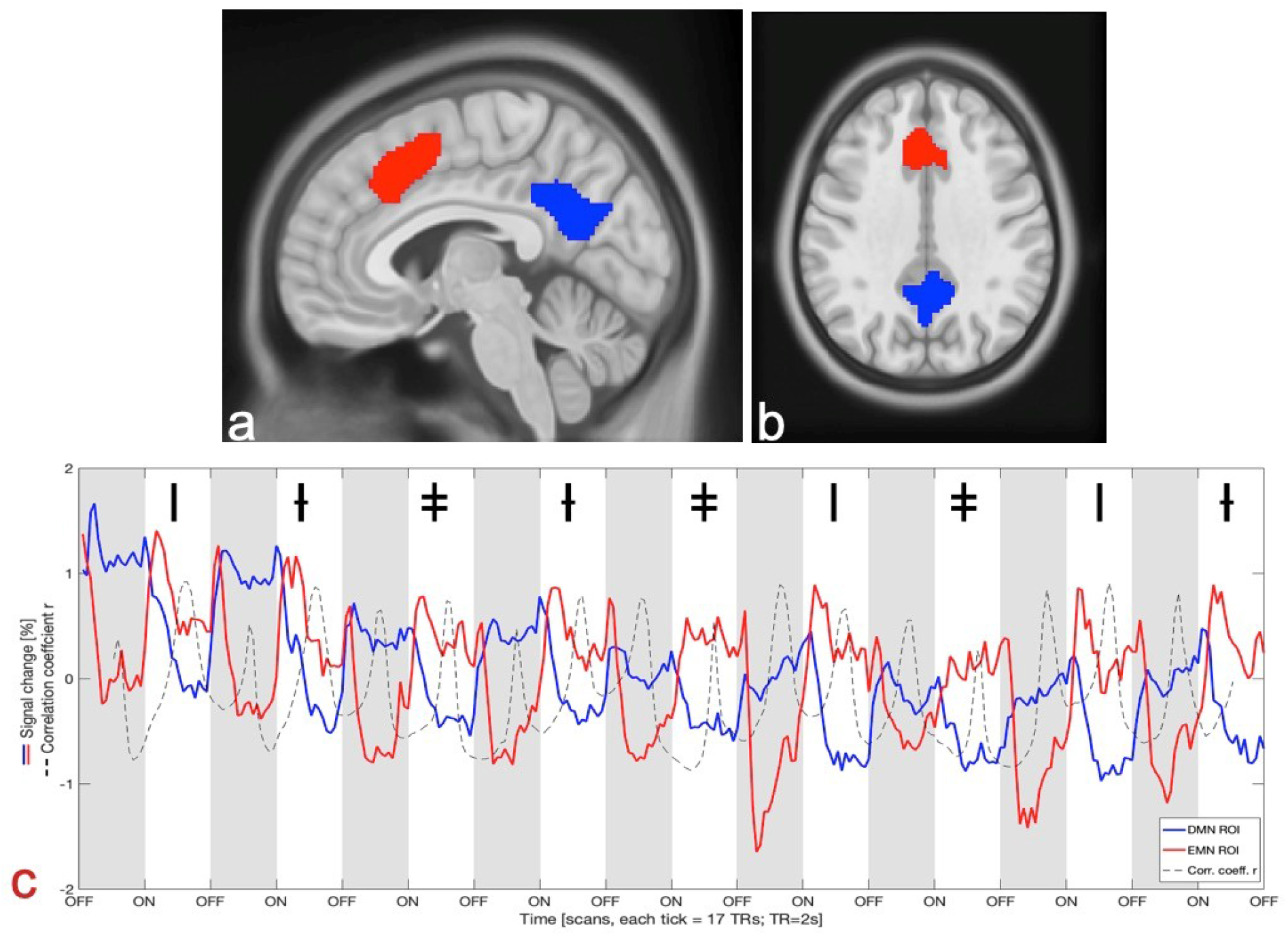
Sagittal (**a**) and axial (**b**) views of the ROIs used for time-series extraction (**c**) shown on the MNI-T1 template. The DMN ROI is shown in blue and the EMN ROI in red. Extracted mean time-series signal change in percent (%) shown in (**c**): Onsets and offsets of the tasks and rest periods are shown on the x-axis as ON- and OFF-blocks, respectively. OFF-blocks are marked in grey. The tasks alternated randomly between mental rotation (marked with the symbol I), working memory (marked with the symbol Ɨ) and mental arithmetic (marked with the symbol ǂ). The black intermittent line in panel (**c**) shows Pearson correlation coefficients continuously calculated on the mean of a 34 sec sliding-window through the whole session, between the DMN- and EMN-extracted time-course signals.

## Discussion

To sum up and discuss the main findings, switching between resting and active processing periods resulted in two distinct networks that were up- and down-regulated as environmental demands changed from passive to active. In this respect, the present results resembled the results from a similar study where the processing tasks were auditory in nature^19^. The existence of a generalized task-positive network is therefore not dependent on sensory modality. The behavioural results confirmed that the subjects were performing the tasks required throughout the scanning session. A second finding was that the up-regulation of the task-positive network was independent of the specifics of the task, and as seen in Figure 2 and 3, generalized across the tasks. In this respect, the present results are in line with the findings of Fedorenko et al.^15^ and Duncan^16^ and point to the existence of a task non-specific network, which we will label the Extrinsic Mode Network (EMN), following the nomenclature from Hugdahl et al.^17^. Considering that the default mode network (DMN) in essence is a task-negative network^2,5,34,35^ being up-regulated in periods of absence of specific processing demands, it is an intrinsic mode network. We now propose that the brain alternates between an intrinsic and extrinsic mode of function, corresponding to the dominating environmental demands, with the intrinsic mode network dominating during task-absence, and the extrinsic mode network dominating during task-presence^19,34^. What is new in the present results is that the extrinsic mode network is a *task non-specific* network, in contrast to *task-specific* task-positive networks, like the salience network^10,11,36^, dorsal attention network^3,37^, or central executive network^12,38^. Thus, the intrinsic and extrinsic mode networks operate at a superordinate level with regard to task-specific networks. We focused on the precuneus/posterior cingulate cortex (PCC) and ventral SMA as ROIs when extracting the time-courses representing the extrinsic and intrinsic mode networks, respectively, since the SMA was overlapping in activation for all three cognitive tasks, and has previously been implicated in a variety of cognitive operations, including visuo-spatial processing, working memory, and mental arithmetic^39^. The precuneus region was chosen because it was strongly activated during OFF-blocks, and has previously been shown to be implicated in the default mode network^1,11,40,41^. Thus, the time-course dynamics seen in Figure 3 from the SMA and precuneus ROIs is taken as a proxy for the up- and down-regulation of the corresponding extrinsic and intrinsic mode networks. We suggest that the role of the SMA, and in particular the ventral portion, is overlapping with the pre-SMA. The conjunction analysis revealed that also other areas activated during ON-blocks were found for the angular gyrus, superior parietal lobule, anterior insula, and middle frontal gyrus. All these regions are implicated in higher-order cognition, including mental arithmetic, visuo-spatial processing, mental rotation and working memory^27,42–44^. In addition, the occipital activation which most likely reflects the fact that the tasks were all visual in nature. Likewise, the activation bilaterally in the pre-central region would be related to the motor-response and button-press. The OFF-blocks were characterized by activations in the angular gyrus and central operculum, which are regions previously implicated in the DMN^32,45^. Looking at Figure 2, which shows area with overlapping and non-overlapping activations for the three tasks reveals a remarkable similarity in activation patterns across all three tasks during ON-blocks. This is evident for all activations, except for the left inferior parietal lobule, which was more strongly activated to the mental rotation and working memory tasks than to the mental arithmetic task. A similar pattern emerged from looking at the overlapping activations for OFF-blocks, i.e. whether activations during resting periods associated with the three tasks differed. Again, as for ON-block periods, the pattern of activations for the OFF-blocks were quite similar across resting periods, especially for the precuneus with centre of gravity showing similar overlapping activations. Activation related to the mental rotation task did however extend in the inferior axis, and was also seen with a small area in the medial-ventral frontal lobe. The patterns of overlapping activations differed however for ON- and OFF-blocks when it comes to overlapping areas for only two tasks. For ON-blocks, the mental rotation and working memory tasks showed overlapping activations (shown in violet colour in Figure 2) to a greater extent than the other task-combinations, which was not the case for OFF-blocks. For OFF-blocks, most of the activations were overlapping for all three tasks, with mental arithmetic (shown in green colour) in addition activating the middle temporal and superior frontal gyri.

These areas are implicated in working memory tasks^46^ and may be overshootings from the ON-periods for this particular task. The ON-OFF contrast resulted in unique activations in the thalamus for mental rotation and mental arithmetic tasks, as well as in left lateral superior frontal gyrus. Thalamus activations have previously been reported for both mental arithmetic and mental rotation^47,48^ and may be related to the role of basal ganglia in number processing and visuo-spatial processing in general. In conclusion, the present results have shown the existence of a generalized task-non-specific network, which is upregulated during periods of active task-processing, but is essentially independent of the specifics of the task. By following the nomenclature introduced by Hugdahl et al.^17^ we have labelled this network the Extrinsic Mode Network (EMN), as an extrinsic mode network in contrast to the default mode network, which is an intrinsic mode network, and should perhaps also be so labelled (IMN). Second, building on our previous research with auditory tasks^19^, the current results have shown that similar network dynamics exist for visual tasks, which in isolation has been shown previously^1,18^. We now show that this also occurs across different tasks in a single scanning session. For these reasons, we now suggest that a standard, and classic, fMRI block-design, with alternating task- and rest-periods, may be an experimental proxy for alternating rest-periods combined with task processing requirement periods that we encounter in the course of an ordinary day in life. Having said that, it should be noted that such periods in reality will vary with regard to intensity, frequency and length as environmental demands vary, and will also vary between individuals.

## Methods

### Subjects

The subjects were 47 adult healthy (from self-report) individuals, 27 males and 20 females, mean age 34.9 years (SD 13.9). The subjects were recruited in the Bergen city area through general, open announcements.

### Stimuli and design

There were three different cognitive tasks and stimuli, a mental arithmetic task, working memory task, and a mental rotation task that were each repeated three times in 30 sec ON-blocks in a random sequence, alternated with 30 sec OFF-blocks (Figure 4).

**Figure 4.**
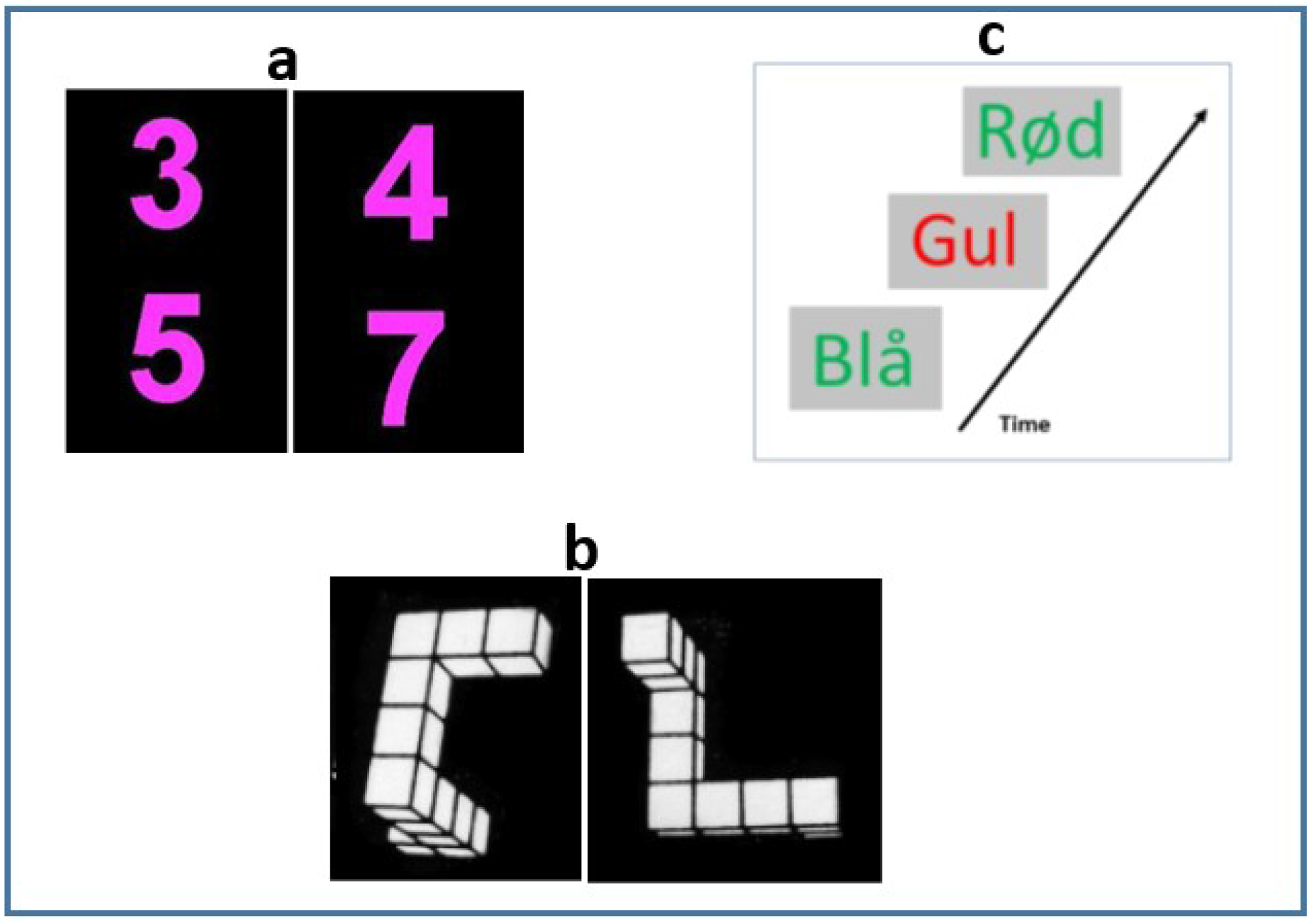
Examples of the stimuli used in the three cognitive tasks. (**a**) Shows examples of the digit pairs shown on each trial, to the left is shown two digits that do not match the target since the sum of the two is < 11. To the right is shown two numbers that match the target since the sum is 11, and the subject should then correspondingly press the response button. (**b**) This shows two of the 3D objects that should be compared for similarity, the example shows two object shapes that are the same but rotated horizontally with regard to each other. The subject should press the response button to this example. (**c**) This shows an example of the 2-back Stroop working memory task in Norwegian, where the instruction is to press the button when the colour of the word matches with colour shown two items before.

The stimuli were constructed with graphics software and stored as images for presentation through high-resolution LCD HD goggles (NordicNeuroLab Inc., Norway, https://nordicneurolab.com/) mounted to the MR head-coil. The timing, duration and sequencing of the stimuli, as well control of overall timing parameters of the experiment was done in the E-Prime (Psychology Software Tools, Inc., USA, https://pstnet.com/) software platform. Synchronization timing of presentation of the stimuli with acquisition of the MR data was done through a NordicNeurolab SyncBox (NordicNeuroLab Inc., Norway, https://nordicneurolab.com/). There were nine ON-blocks with task presentations (three repetitions of each task), alternated with nine black screen OFF-blocks without task-presentations. The OFF-blocks had a fixation-cross in the middle of the visual field, and the subjects were instructed to keep their eyes open during OFF-blocks. Each ON- and OFF-block lasted for 60 sec, making a total of 18 min.

The mental arithmetic task consisted of two digits that could vary between the digits 1-9, written on top of each other, in purple font against black background (see Figure 4(a) for examples). Each pair of digits was presented for 300 ms, with 20 pseudo-random trials in each ON-block. The instruction to the subject was to mentally add the two numbers and press a hand-held button whenever the sum was “11”. There were five target trials randomly interspersed among the 20 trials (40%), with the restriction of not allowing two or more consecutive targets by random selection. Thus, there were 15 target trials in total.

The mental rotation task (Figure 4(b)) was tailored on the classic Shephard and Metzler mental rotation task^24^ with images of two 3D non-configurative objects in white against black background, that were displayed on each trial. The task of the subject was to decide by pressing the hand-held button whether the two shapes were two different objects, or two shapes of the same object, but where they were rotated horizontally with regard to each other, and where the rotation varied between 20° and 180°. Each presentation trial lasted for 300 ms, with 20 trial presentations during an ON-block, and where the two shapes were the same on 10 trial presentations and different on 10 trial presentations. The instruction was to press on “same”, thus there were a total of 30 targets across the Mental rotation task.

The working memory task (Figure 4(c)) was an n-back task with presentations of incongruent Stroop colour-words, like the word “red” written in a blue ink. The instruction to the subject was to remember the colour of the word presented two items back (2-back paradigm), and to press the hand-held when the item currently seen in the goggles matched the one presented two items back. This task adds an executive function process to the working memory process, making it cognitively more demanding. It has previously been shown to discriminate between healthy and depressed patients^28^, and reliably elicits demands for working memory processing. Each presentation lasted for 300 ms, with 20 presentations in total, and with five (40%) pseudo-randomly interspersed target trials where the two displays matched. Thus, there were a total of 15 targets for the working memory task.

#### Behavioural data statistics

Behavioural data for the three tasks were available for 46 participants, one missing data set. were automatically collected and stored by the E-prime software, for later statistical analysis of response accuracy and response latency.

For all three tasks, response accuracy, and response latency (RT ms) was evaluated statistically for means, standard deviations, and hits ratio of correct responses to false alarms, respectively. Hits ratios were converted to Cohen's d-statistic for effect sizes for each task.

### Procedure

The subjects were first interviewed for body implants, like pace-makers, and any signs of claustrophobia, and were then presented with the informed consent form which they signed. Females were in addition asked about pregnancy, which was an exclusion criterion. The study was approved by the Regional Committee for Medical Research Ethics in Western Norway (2014/1641/REK Vest). They were then informed about the study and shown examples of the stimuli and familiarized with the tasks to be presented when in the scanner, before being placed in the scanner. To facilitate direct communication with the MR-technicians in the control-room, a balloon was placed on the chest, which should be squeezed in case of an emergency. There was also direct audio contact between the scanner chamber and the control room. The E-Prime stimulus program was run from a stand-alone PC and operated by a research assistant.

#### MR scanning and data acquisition

The experiment was conducted on a 3T Magnetom Prisma MR scanner (Siemens Healthcare, Germany). An anatomical T1-weighted image was acquired prior to the functional imaging with the following sequence parameters: MPRAGE 3D T1-weighted sagittal volume, TE/TR/TI = 2.28ms/1.8s/900ms, acquisition matrix = 256×256×192, field of view (FOV) = 256×256mm2, 200 Hz/px readout bandwidth, flip angle = 8 degrees and total acquisition duration of 7.40 minutes. 2D gradient echo planar imaging (EPI) was performed with the following parameters: TE/TR = 30ms/2s, 306 volumes in time, acquisition matrix = 64×64, slice thickness = 3.6 mm, 35 slices, FOV = 230×230mm2.

#### fMRI data processing

All pre-processing and analysis were performed in Matlab 9.5 (the MathWorks, Natick, MA) using SPM12-r7219 (the Wellcome Centre for Human Neuroimaging, UCL, London, UK, https://www.fil.ion.ucl.ac.uk/spm/). Pre-processing consisted of rigid-realignment, normalization to MNI space and smoothing with a 8×8×8 mm3 Gaussian kernel.

#### Conjunction Analysis

After pre-processing, first level analysis for all subjects was performed. The resultant contrast images were then used to perform a second level conjunction (one-way ANOVA) analysis across all OFF>ON and ON>OFF images of the individual tasks. Statistical maps were calculated with correction for multiple comparisons (family-wise errors, FWE), a *p*-value of 0.05 and cluster extent of 20 voxels.

The tasks were also studied individually adjunct to the conjunction analysis to assess the effect of each task. These individual maps were processed in MRICron (ch2.better.niftii) and superimposed on each other.

#### ROI time course analyses

A third analysis concerned comparing and contrasting activity in regions of interest (ROI) that were functionally defined from the activation patterns seen in the conjunction analyses, see Figure 3. The ROIs were chosen because these regions have shown significant activations during resting- and task-absent periods^4,5,32,49^. Similarly, they have shown ROI-activations during active processing and task-presence^15,19,28,29,50^ including tasks and processes covered by the current three tasks.

For this time-course analysis, mean voxel values over the ROI were extracted for each subject and averaged over the cohort. Resultant time-course curves were normalized and rescaled as a percentage of total BOLD signal change. ROI were created using MarsBaR v0.44^51^.

## Acknowledgments

The present study was funded by grants to R.G. from the Trond Mohn Foundation (#BFS2017TMT06) and to K.H. from the European Research Council (#693124) and the Health Authorities of Western Norway Helse-Vest #912045).

The authors want to thank all subjects participating in the study, and the MR technicians and research technicians who assisted in acquiring the data.

## Author Contributions

F.R. carried out analysis of the data, manuscript writing and figure preparation. R.G. conceived and conducted the experiments. J.B. developed the fMRI tasks and analysed response data.

K.K. contributed to analysis of the data. L.E. conceived and conducted the experiments. K.H. conceived and conducted the experiments and carried out analysis of the data. All authors reviewed the manuscript.

## Additional Information

Correspondence and requests for materials should be addressed to F.R.

